# Interleukin-6 Signaling Mediates Cartilage Degradation and Pain in Post-Traumatic Osteoarthritis

**DOI:** 10.1101/2021.09.08.459303

**Authors:** Yihan Liao, Yinshi Ren, Xin Luo, Jason T. Long, Anthony J. Mirando, Abigail P. Leinroth, Ru-Rong Ji, Matthew J. Hilton

## Abstract

Osteoarthritis (OA) and post-traumatic OA (PTOA) are prevalent joint disorders and leading causes of chronic pain. The disease pathology of OA/PTOA is caused by imbalanced catabolic and anabolic responses and pro-inflammatory changes; however, their connection to pain is not well studied. Since IL-6 is involved in cartilage degradation and conditions of inflammatory pain, we set out to identify whether IL-6 and IL-6 signaling mechanisms contribute to both PTOA-associated cartilage degradation and pain. We performed a modified destabilization of the medial meniscus (DMM) surgery, a model of PTOA, on conventional IL-6 KO and control mice and assessed both cartilage degradation and pain-associated phenotypes. Genetic removal of *Il6* in males attenuates PTOA-associated cartilage catabolism, decreases innervation of soft tissues associated with the knee joint, and reduces nociceptive pain signaling, without improving subchondral bone sclerosis or chondrocyte apoptosis. We further demonstrate that specific downstream mediators of IL-6 signaling, the Janus kinases (JAKs), are critical in regulating both cartilage catabolism and pain signaling. We identified STAT3 as a key regulator of cartilage catabolism downstream of JAK; however, inhibition of STAT3 decreases cartilage anabolism while enhancing pain signals. ERK was found to be important for neurite outgrowth and pain signaling; however, inhibition of ERK was less effective in reducing cartilage catabolism. Therefore, our data demonstrate that IL-6 mediates both PTOA-associated cartilage degradation and pain, and provides critical details regarding the downstream mediators of IL-6 signaling as therapeutic targets for disease-modifying osteoarthritis drugs.

Single Sentence Summary

IL-6 mediates PTOA-associated cartilage degradation and pain via specific downstream signaling mechanisms in a gender specific manner.

## Introduction

Osteoarthritis (OA) is the most prevalent joint disorder and a leading cause of disability, contributed by multiple risk factors, including aging, increased body mass index (BMI), and prior joint injury among others^1–3^. Post-traumatic OA (PTOA), or injury induced OA, accounts for approximately 12% of symptomatic OA^4^. Pain is a dominant symptom of OA/PTOA and serves as a key criteria in the diagnosis and treatment^5^. OA/PTOA is a major contributor to chronic pain (~35%) affecting nearly 32.5 million people (1 in every 10 adults), which translates to an estimated $65.5 billion of annual medical costs in the US alone^6^. Despite the extensive socioeconomic burden of OA/PTOA, no effective disease-modifying osteoarthritis drugs (DMOADs) are yet available for OA/PTOA-associated joint degeneration and pain management. Current treatments, such as topical capsaicin, acetaminophen, NSAIDs, and corticosteroid injections, either have limited efficacy or variable side effects^7, 8^.

OA/PTOA is not simply a disease affecting structural alterations to the articular cartilage, but rather a disease affecting the entire joint, which can involve the ligaments, meniscus, nerves, subchondral bone, synovial membrane, and periarticular muscles^9, 10^. The physical manifestations of OA/PTOA, such as articular cartilage and meniscus degeneration, synovial hyperplasia, osteophyte formation, and subchondral bone sclerosis, are caused by an imbalance in anabolic and catabolic responses, as well as, pro-inflammatory changes^11, 12^. Factors involved in OA/PTOA initiation and progression include pro-inflammatory cytokines (TNFα, IL-1β, IL-6, and IL-17) and matrix degrading enzymes (MMP-1, MMP-3, MMP-9, MMP-13, ADAMTS-4, ADAMTS-5)^13^. In addition to structural changes to the joint, the pathology of OA/PTOA-associated pain has also been under investigation. As in all types of chronic pain, OA/PTOA-associated pain is the dynamic result of a complex interaction between local tissue damage and inflammation, peripheral and central sensitization, and the brain^14^. Tumor Necrosis Factor Alpha (TNFα), Interleukin-1 Beta (IL-1β), Interleukin-6 (IL-6)^15–17^, C-C Motif Chemokine Ligand 2 (CCL2)^18, 19^, and the neurotrophic factor, Nerve Growth Factor (NGF),^20, 21^ are all reported as being correlated with OA/PTOA-associated pain. Since IL-6 is both a major predictor of OA/PTOA^22^ and has been associated with inflammatory pain in a number of contexts^23^, it may serve as a critical effector connecting OA/PTOA-associated joint changes and pain.

IL-6, as an inflammatory cytokine, has been implicated in both OA/PTOA cartilage degeneration and other aspects of inflammatory pain. IL-6 signals by binding to the IL-6 specific membrane receptor (mIL-6R), known as ‘classical signaling’, or by binding to the soluble form of IL-6R (sIL-6R), known as ‘trans-signaling’. Both interactions activate the transducer glycoprotein 130 (gp130) and induce a cascade of Janus Kinase (JAK) protein phosphorylation, which further leads to the phosphorylation and activation of Signal Transducer and Activator of Transcription (STAT) and/or Mitogen-Activated Protein Kinase (MAPK) proteins ^24, 25^. Elevated levels of IL-6 have been identified in both the articular cartilage^26^ and synovial fluid^27^ in human OA cohort studies, while a large scale, long-term follow up (15-years) study identified elevated levels of serum IL-6 as a significant predictor of radiographic knee OA^22^. In murine OA studies, intraarticularly injecting recombinant IL-6 induced cartilage destruction after 14 days and DMM-surgery, a joint injury model of PTOA^28, 29^, induced less cartilage destruction in IL-6 knockout mice^30^. Our previous work demonstrated that p-STAT3, a downstream mediator of IL-6 signaling, is elevated within cartilage and synovium following DMM-induced PTOA^31^. Other work has indicated that the STAT3 inhibitor, Stattic, results in reduced cartilage degradation during DMM-induced PTOA in vivo^32^. In addition to its role as a cytokine during immune responses, IL-6 signaling also plays a role in the nervous system where IL-6 and IL-6R are expressed in sympathetic and sensory ganglia^33^. IL-6 has been linked to chronic pain through the regulation of the pain-activated ion channel TRPV1^34, 35^, and is known to induce neurite outgrowth of the PC12 neuronal cell line *in vitro*, similar to effects caused by NGF.^33, 36^ Based on these evidences, we hypothesized that IL-6 may not only be involved in PTOA-associated cartilage degeneration, but also PTOA-associated pain.

Although accumulated evidence indicates a contribution of IL-6 to cartilage degradation in OA/PTOA progression, no direct link between IL-6 and PTOA-associated pain has been established. Here, we utilize the DMM surgical model of PTOA with conventional IL-6 knockout (KO) and control mice to study the function and mechanisms of IL-6 signaling in both cartilage degeneration and pain signaling within nociceptive neurons associated with the knee joint. Our findings demonstrate that IL-6 mediates both PTOA-associated cartilage degradation and pain in males, and indicate specific roles for downstream mediators of IL-6 signaling within chondrocytes and nociceptive neurons, which have important implications for therapeutic targets of disease-modifying osteoarthritis drugs.

## Results

### Genetic removal of *Il6* alleviates PTOA-associated cartilage catabolism

To determine the necessity of IL-6 in PTOA-associated cartilage degeneration and pain, we performed both sham and a modified DMM surgery on conventional IL-6 KO (*Il6^−/−^*) and wild-type (WT) control mice at 4-months of age, followed by assessments of joint cartilage degeneration and pain at 4-, 8- and 12-weeks post-injury (wpi). Similar to prior studies, our modified DMM surgery induced a progressive OA phenotype in wild-type (WT) mice as shown via histological assessments and confirmed with OARSI scoring from 4-weeks to 12-weeks following DMM injury (Supplementary Figure 1). To identify any innate differences between the two mouse strains, we characterized WT and *Il6^−/−^* mice with sham injury first, and didn’t detect any significant difference in cartilage, subchondral bone, knee innervation, or pain sensation at either the molecular or behavioral level (Supplementary Figure 2). These data allow us to compare the two mouse strains directly in the following experiments.

In *Il6^−/−^* mice, DMM surgery induced minor OA phenotypes with a reduction in Safranin-O staining and/or small fibrillations at the cartilage surface, similar to that seen in WT mice at 4-weeks post-DMM (Figure 1A). However, at 8- and 12-weeks after DMM injury, *Il6^−/−^* mice show significantly less cartilage degradation than WT mice, which exhibit severe clefting and cartilage erosions at both time points. This is evidenced by the significantly lower OARSI scores in *Il6^−/−^* mice at 8- and 12-weeks post-DMM when compared to WT controls (Figure 1B,1C). Further, OARSI scores of *Il6^−/−^* mice following DMM were noted as similar to WT sham controls at these later timepoints, indicating that genetic removal of *Il6* is capable of attenuating cartilage degradation in PTOA for prolonged periods of time.

**Figure 1.**
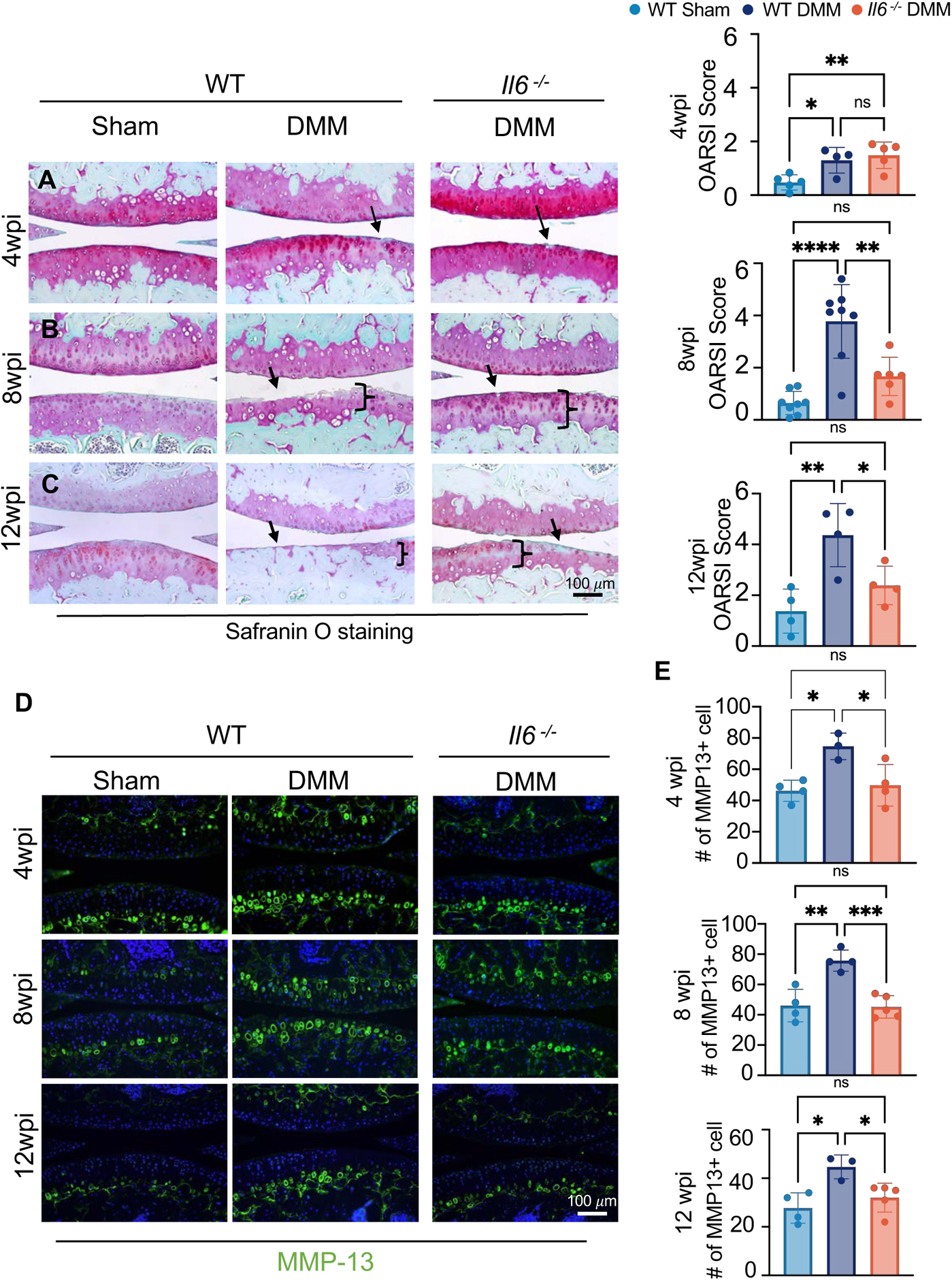
Loss of IL-6 results in the reduction of cartilage degradation in PTOA. (A) Safranin-O staining of WT and *Il6−/−* knee joint sections at 4- (A), 8- (B) and 12- (C) weeks post injury in male mice. Bar = 100 μm. On the right, OARSI score quantified the cartilage phenotype at 4- (N ≥ 4), 8- (N≥ 6), and 12- (N≥ 4) weeks post injury. p<0.05; one-way ANOVA. (D) IF staining of MMP-13 (green) of knee sections from WT or *Il6−/−* mice at 4-, 8- and 12-weeks post sham or DMM surgery. Bar = 100 μm. (E) Quantification of the number of cells with MMP-13 staining. N≥4. p<0.05; One-way ANOVA. Data represent mean ± SD.

In PTOA, articular cartilage degradation is generally contributed to by the imbalance of cartilage anabolism/catabolism (remodeling) and chondrocyte cell death. To understand how removal of *Il6* affects cartilage degradation in PTOA, we first examined levels of the catabolic protein, MMP-13, using immunofluorescent (IF) staining on knee joint sections from *Il6^−/−^* and WT control mice at 4-, 8-, and 12-weeks post-DMM. An upregulation of MMP-13 levels were induced by DMM in WT mice at all time points, but there were significantly less MMP-13 positive cells in *Il6^−/−^* joint chondrocytes as compared WT mice, indicating a reduction in catabolism of *Il6^−/−^* mice from early to late timepoints following DMM injury (Figure 1D, 1E). Similar to the OARSI scores, MMP-13 levels in *Il6^−/−^* joint chondrocytes following DMM were comparable to WT sham controls. To examine whether the reduced cartilage degradation observed in *Il6^−/−^* knee joints following DMM occurs via altered chondrocyte survival, we performed TUNEL staining on knee joint sections from *Il6^−/−^* and WT control mice at 4-, 8-, and 12-weeks post-DMM. We observed that at 4-weeks post-DMM, in both WT and *Il6^−/−^* mice, there is a minor increase in cell apoptosis within the articular cartilage after injury (Supplementary Figure 3A); however, by 8-weeks post-injury there were significantly increased cell apoptosis in the superficial and deep zones of both WT and *Il6^−/−^* mice as compared to sham control. Importantly, we observed no difference between the DMM injury groups of WT and *Il6^−/−^* mice (Supplementary Figure 3B). Collectively, these observations indicate that IL-6 regulates cartilage degradation in PTOA via the control of MMP-13-mediated cartilage catabolism and not by altering chondrocyte cell death.

### The loss of IL-6 results in the reduction of pain in PTOA

Since pain is a prominent symptom of OA and IL-6 is known to be involved in various settings of pain, we measured local knee pain in both WT and *Il6^−/−^* mice following DMM/sham injury utilizing a pressure application measurement (PAM) device. The PAM device measures the precise max force that a mouse can withstand at the knee joint prior to vocalization and/or limb withdrawal^37, 38^. The lower the paw withdrawal threshold, the higher the pain sensitivity. In WT mice, we observed at 1-week post-injury, there is significant pain caused by the surgery in both sham and DMM injured mice. In the sham group, surgical pain recovers over time, and returns to baseline by 8-weeks post-surgery (Figure 2A). On the contrary, local knee pain in WT DMM mice is significantly enhanced when compared to sham from 4- to 8-weeks post-injury, indicating DMM injury induces chronic knee pain in WT mice (Figure 2A). However, *Il6^−/−^* mice with DMM injury exhibit significantly less pain from 2- to 8-weeks post-injury when compared with WT DMM injured mice and demonstrate a complete recovery and return to baseline levels over this time (Figure 2A).

**Figure 2.**
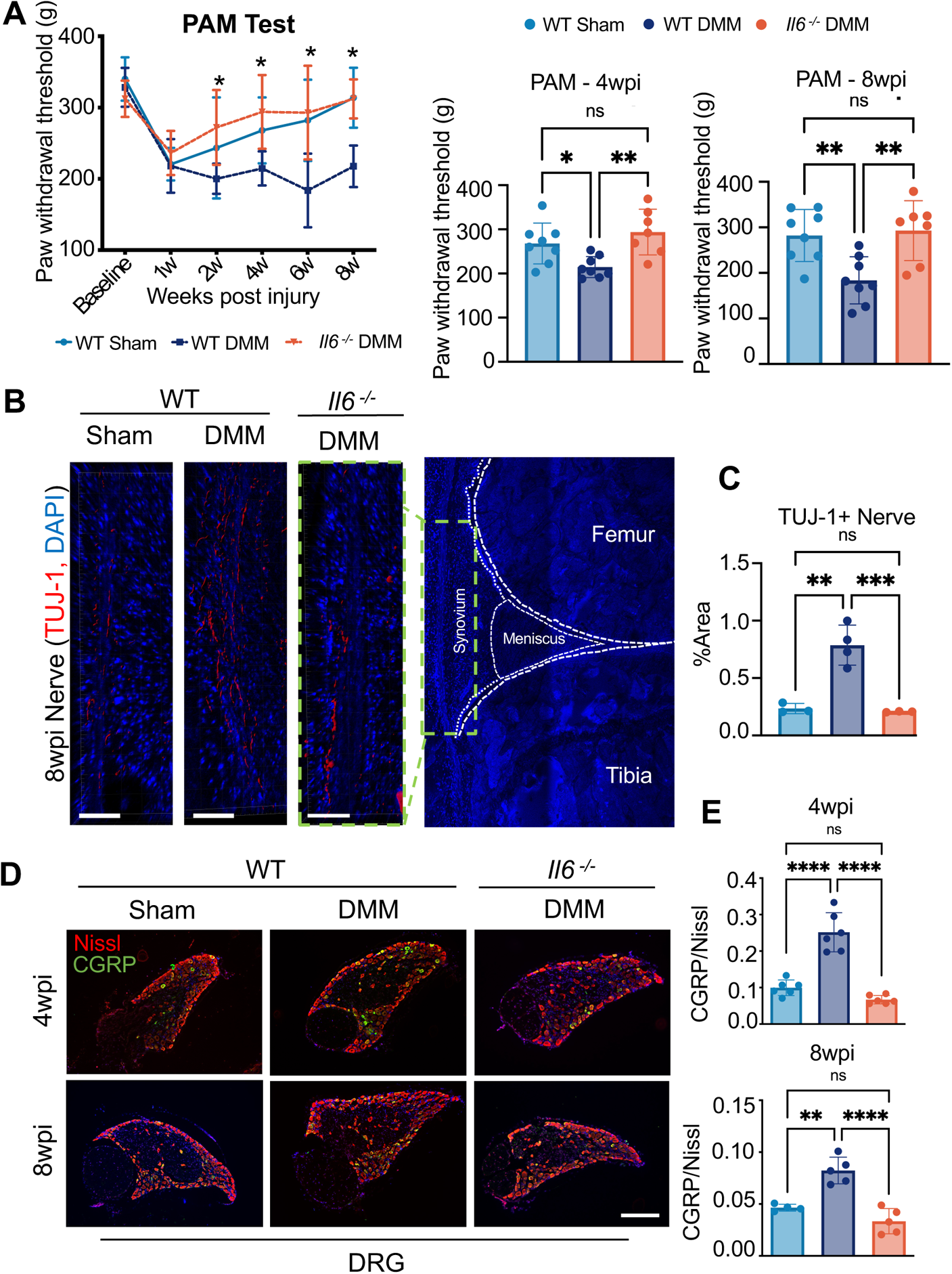
Loss of IL-6 results in the reduction of pain in PTOA. (A) Behavioral tests (PAM) for knee pain (mechanical hyperalgesia) in male mice. Paw withdrawal threshold was measured with a PAM device when pressure applied to the knee joint. N≥ 8, p<0.05; Two-way ANOVA with Bonferroni’s post-hoc test. * indicates the statistical significance between control DMM and IL-6 KO DMM group. (B-C) IF staining and its quantification showing TUJ-1+ innervation at knee. TUJ-1 in red. N≥ 3 (D) IF of CGRP+ neurons in L3-5 DRG from WT or *Il6^−/−^* mice following sham or DMM surgery at 4- and 8-weeks post injury. CGRP in green, Nissl in red. Bar = 200 μm (E) Quantification of CGRP/Nissl neurons. N≥ 4 p<0.05; One-way ANOVA. Data represent mean ± SD.

To investigate the underlying mechanisms that may account for the reduced pain response observed in *Il6^−/−^* mice following DMM/sham injury, we first characterized innervation of the sham and DMM-injured knee joints of WT and *Il6−/−* mice by immunostaining joint sections for TUJ-1. We observed an enhancement of innervated synovial tissues within WT mice following DMM injury, but not in *Il6^−/−^* mice (Figure 2B, C). To assess their molecular response to pain, we next measured levels of the pain-associated molecules, Calcitonin Gene Related Peptide (CGRP) within specific dorsal root ganglia (DRG). The DRG are a cluster of nociceptive sensory neurons that transmit pain signals from the peripheral to central nervous system. Specifically, Lumbar 3-5 (L3-L5) DRG correspond to the peripheral nerves innervating knee joint tissues, including the synovium, ligaments, osteochondral junction, and meniscus. By performing Nissl staining on DRG tissue sections we can visualize all neurons within a DRG, while incorporating IF staining for CGRP can provide a ratio of pain sensing neurons within those DRG. A higher ratio of CGRP+ cells (green/yellow) are observed in the DRG of WT mice following DMM injury as compared to WT sham mice at 4- and 8-weeks following surgery. While in *Il6^−/−^* DRG sections, CGRP+ cell ratios in DMM groups were similar to that of sham at both early (4-weeks post-injury) and late timepoints (8-weeks post-injury) (Figure 2D, E). These data are consistent with the PAM behavioral pain tests, suggesting an early rescue of the pain response when deleting *Il6* in mice with DMM injury. Interestingly, pain measures, including both behavioral and molecular readouts on DRG sections, were rescued in *Il6^−/−^* mice at a timepoint when cartilage damage is minimal and remains comparable between in *Il6^−/−^* and WT control mice following DMM injury. These data suggest a potential uncoupling of pain and overt cartilage degradation in PTOA, however IL-6 appears to play a significant role in both PTOA-associated cartilage degeneration and pain.

Since OA is a disease of the whole joint and OA progression and pain has been associated with aberrant subchondral bone remodeling and innervation^39, 40^, we utilized uCT to evaluate the subchondral bone in WT and *Il6^−/−^* knees following DMM/sham surgeries. At 8-weeks post-DMM injury, we observed subchondral bone sclerosis in both WT and *Il6^−/−^* mice (Figure 3), suggesting that removal of *Il6* does not attenuate subchondral sclerosis, and that the reduced PTOA-associated pain and cartilage degradation in *Il6^−/−^* mice does not occur due to corrections in subchondral bone remodeling.

**Figure 3.**
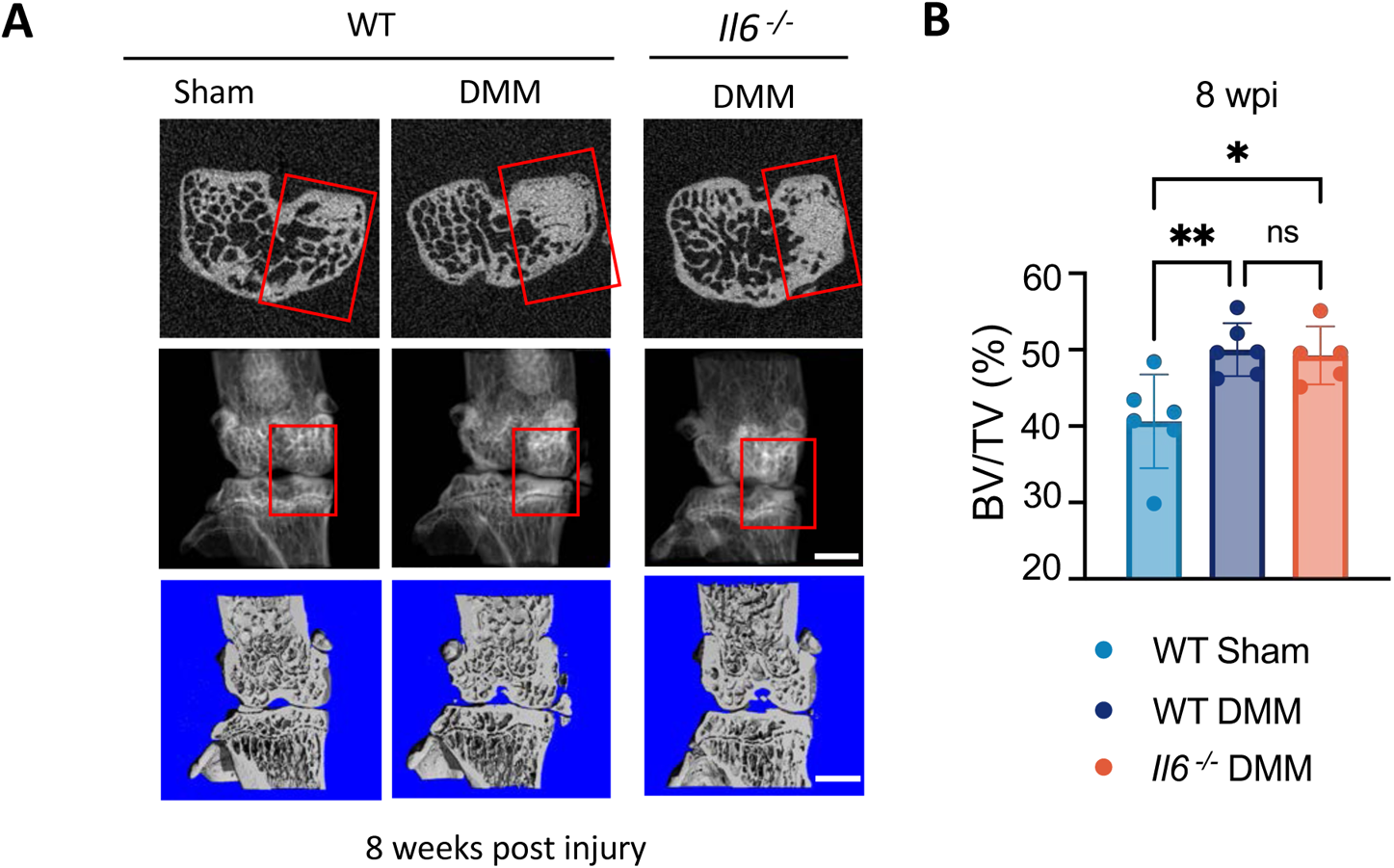
MicroCT evaluation of OA phenotypes within subchondral bone. (A) MicroCT analysis of WT or *Il6−/−* mice following sham or DMM surgery at 8-weeks post injury. Bar = 1 mm. (B) Quantification of subchondral bone with 0.1mm thickness of the region of interest is evaluated. WT N=6; KO N=5, p<0.05; One-way ANOVA. Data represent mean ± SD.

### *Il6* deletion in female mice does not affect PTOA-associated cartilage degradation and pain

In addition to the results evaluated in male mice as described above, we also examined female joint cartilage and pain phenotypes associated with PTOA. As previously shown, female mice exhibit a slower and blunted cartilage catabolic response to DMM injury as compared to male mice^41^. We did not observe significant cartilage degradation until 12-weeks post-injury, which was still only highlighted by small cartilage fibrillations and vertical clefts at the surface of the cartilage producing an OARSI score of 2 on average. Histology and OARSI scores of *Il6^−/−^* female knee sections indicate a similar PTOA cartilage phenotype as compared to WT female mice following DMM injury, suggesting that removal of *Il6* in females does not provide significant chondroprotection at this stage (Supplementary Figure 4A, 4B). As for pain, both WT and *Il6^−/−^* female mice exhibit persistent knee pain (Supplementary Figure 4C). At 12-weeks post-DMM, when the PTOA-associated cartilage phenotypes were visible, *Il6^−/−^* female mice continued to show significant knee pain when compared with sham female mice (Supplementary Figure 4D). The failure to rescue PTOA-associated cartilage degradation and pain in female mice via the genetic removal of *Il6* suggests that at least some of the molecular mechanisms regulating PTOA-associated cartilage degeneration and pain differ between males and females.

**Figure 4.**
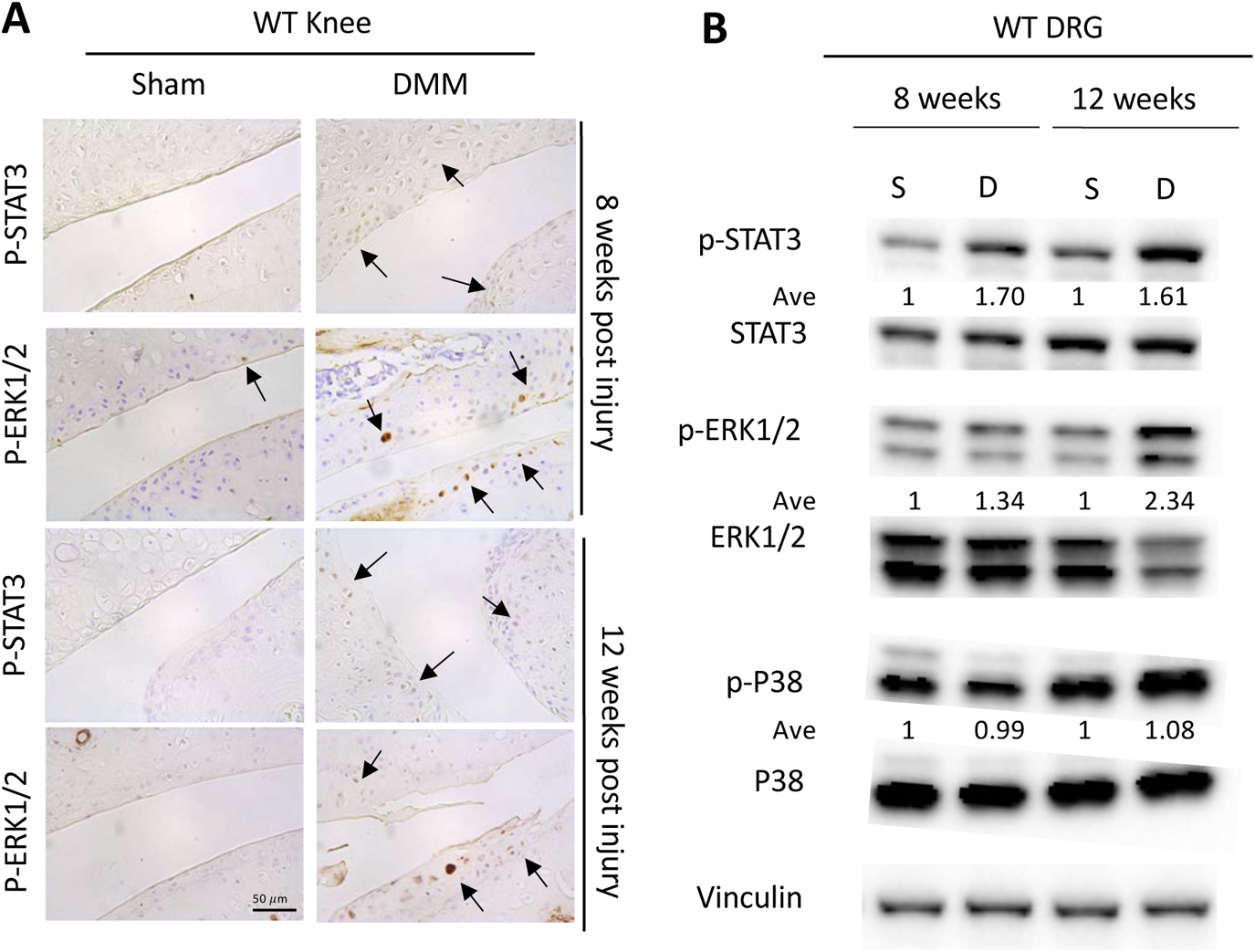
IL-6 signaling within joint cartilage and DRGs following DMM injury. (A) IL-6 downstream signaling p-STAT3 and p-ERK IHC in knee of WT mice following sham or DMM surgery at 8- and 12-weeks post injury. 8wpi N≥ 4; 12wpi N≥ 4. Bar = 50 μm. (B) Western blot showing IL-6 downstream signaling in L3-5 DRG from WT mice following sham or DMM surgery at 8- and 12-weeks post injury. 8wpi N≥ 3; 12wpi N≥ 5.

### Multiple downstream mediators of IL-6 signaling are upregulated in joint tissues and peripheral nerves following joint injury

To further understand the molecular mechanisms by which IL-6 signaling may regulate cartilage degradation and pain in PTOA, we examined the change of downstream mediators of IL-6 signaling in both cartilage and L3-L5 DRG following DMM injury. At 8- and 12-weeks post-DMM injury, we observed a significant upregulation of p-STAT3 and p-ERK within cartilage and synovial tissues (Figure 4A). Distant from the injury site at the knee, DRG associated with L3-L5 vertebrae also revealed an upregulation of p-STAT3 and p-ERK, but not p-P38 at 8- and 12-weeks following DMM (Figure 4B). These data indicate that the activation of downstream IL-6 signaling mechanisms in knee joint tissues and their associated DRG neurons may play essential roles in PTOA-associated cartilage degeneration and pain. Notably, MAP kinase signaling has been strongly implicated in the pathogenesis of pain^42^.

### Specific downstream effectors mediate IL-6 induced cartilage catabolism and pain-associated cellular responses

To better understand the specific effects of IL-6 signaling within cartilage tissue, we first cultured cartilage explants harvested from 3-week-old mouse joints and treated them with recombinant IL-6 protein (rIL-6) *ex vivo.* IL-6 treatment decreased proteoglycan content (Safranin-O staining) and COL2A1 expression (anabolism), while increasing MMP-13 expression (catabolism), all of which contribute to OA-associated cartilage degeneration (Figure 5A). Similarly, we treated ATDC5 chondrogenic cells with rIL-6 for 24 hours and observed a downregulation in the expression of anabolic genes, *Acan and Col2a1*, and an upregulation in the catabolic genes, *Mmp13* and *Adamts5* (Figure 5B). To investigate which downstream mediators of IL-6 signaling regulate these catabolic/anabolic cartilage responses, we utilized specific JAK, STAT, and ERK inhibitors following the treatment of ATDC5 chondrogenic cells with rIL-6 (Figure 5C and Supplementary Figure 5B). We demonstrate that inhibition of IL-6 induced JAK signaling with Ruxolitinib alleviates the IL-6 mediated reduction in *Acan* expression (anabolism), while decreasing the IL-6 mediated induction of *Mmp13* expression (catabolism) (Figure 5C). Inhibition of STAT3 activation utilizing the small molecule inhibitor, Stattic, had similar effects on blocking IL-6 induced *Mmp13* expression, however further suppressed *Acan* expression that is already downregulated by rIL-6 (Figure 5C). Inhibition of ERK1/2 activation utilizing the chemical inhibitor U0126 following rIL-6 treatment did not counteract the IL-6 reduction in anabolic gene regulation, while only having modest effects on the dampening of IL-6 induced *Mmp13* expression (Figure 5C). Based on these *in vitro* experiments, JAK signaling, upstream of STAT3 and ERK signaling, appears to be a key effector in the regulation of IL-6 induced cartilage degeneration through decreasing anabolism and increasing catabolism. While inhibition of STAT3 aids in reducing IL-6 induced cartilage catabolism, the negative effects on cartilage anabolism *(Acan* expression) are worsened. This effect potentially occurs due to an upregulation of pP38, which is not observed following JAK or ERK inhibition (Figure 5C, Supplementary Figure 5B). Collectively, these data further indicate that STAT3 or ERK inhibition alone may not be sufficient to maintain healthy cartilage following the pathologic activation of IL-6 signaling in chondrocytes.

**Figure 5.**
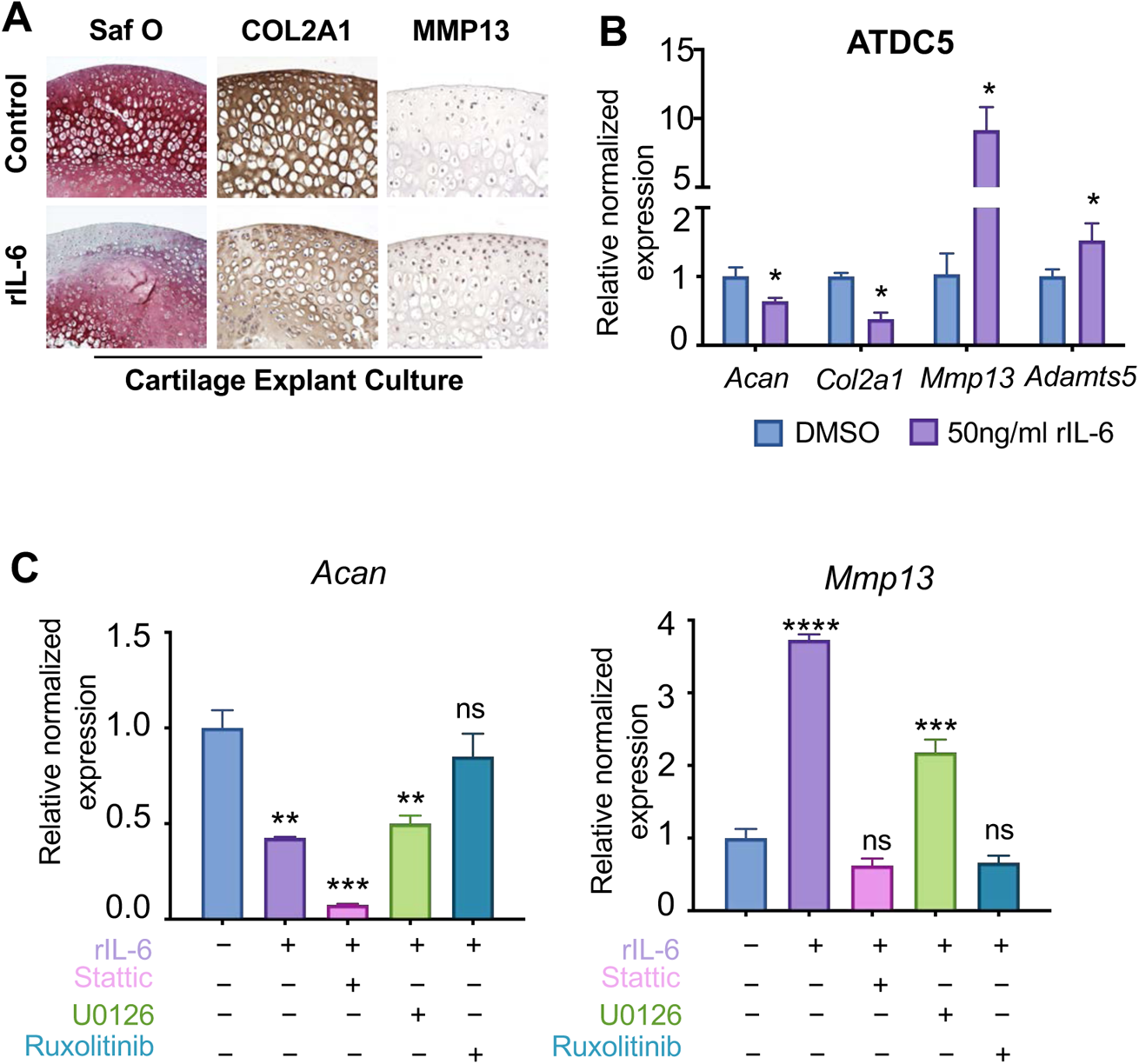
IL-6 induces OA-associated molecular and cellular phenotypes *in vitro* and *ex vivo*. (A) Three-week old mice femoral cartilage explant cultures demonstrate that recombinant IL-6 (rIL6) protein increases catabolic factors and reduces proteoglycan and anabolic factors ex-vivo, N=5. (B) qPCR showing the alterations of gene levels with the induction of 50ng/ml IL-6 in ATDC5 cell cultures at 24-hours. Data represent mean ± SEM. (C) qPCR of ATDC5 cell culture with DMSO or 25ng/ml IL-6 plus or minus inhibitors for 48 hours (0.5uM Ruxolitinib; 20uM Stattic; 10uM UO126). N≥ 3, p<0.05. Data represent mean ± SEM. * indicates statistical significance by one-way ANOVA between DMSO group and treatment group.

To understand the peripheral nerve response to IL-6 signaling and its downstream mediators, we harvested L3-L5 DRGs and cultured as either DRG explants (Figure 6) or primary neuron cultures (Supplementary Figure 6) and treated them with rIL-6 or recombinant NGF (rNGF). NGF is a well-established neurotrophic factor, which can stimulate neurite outgrowth and induce pain related factors such as neuropeptide CGRP^43^. In our DRG explant or isolated primary neuron cultures, neurite outgrowth of DRG neurons were stimulated by rIL-6 in a similar manner to rNGF activation (Figure 6A-B and Supplementary Figure 6). In addition to neurite outgrowth, expression of the pain-associated factors, *Cgrp* and *C-C motif chemokine receptor 2* (*Ccr2*), also exhibited significant upregulation via rIL-6 treatments of DRG explant cultures (Figure 6C). Similar to the ATDC5 chondrocyte cultures, we validated the efficacy of JAK, STAT, and ERK inhibitors and assessed the ability of rIL-6 to induce neurite outgrowth and the expression of specific pain-associated factors in both DRG explant and primary DRG neuron cultures (Figure 6A-C; Supplementary Figures 5C and 6). Again, inhibition of JAK signaling with Ruxolitinib attenuates both neurite outgrowth and pain-associated factors induced by rIL-6. Unlike what we observed in chondrocytes, inhibition of ERK1/2, but not STAT3, results in a rescue of the rIL-6 mediated pathologic effects on DRG neurons and the induction of pain-associated factors. Importantly, treatment with U0126 reduces neurite outgrowth, as well as, the expression of *Cgrp* and *Ccr2* induced by rIL-6, while treatment with the STAT3 inhibitor, Stattic, exhibited minimal effects on neurite outgrowth while exacerbating the rIL-6 induced expression of the pain-associated factors, *Cgrp* and *Ccr2*. These data suggest that inhibition of STAT3 would have no effect or potentially agonistic effects on PTOA-associated pain, while JAK and ERK inhibition may be a more relevant approach to treating IL-6 induced or PTOA-associated pain.

**Figure 6.**
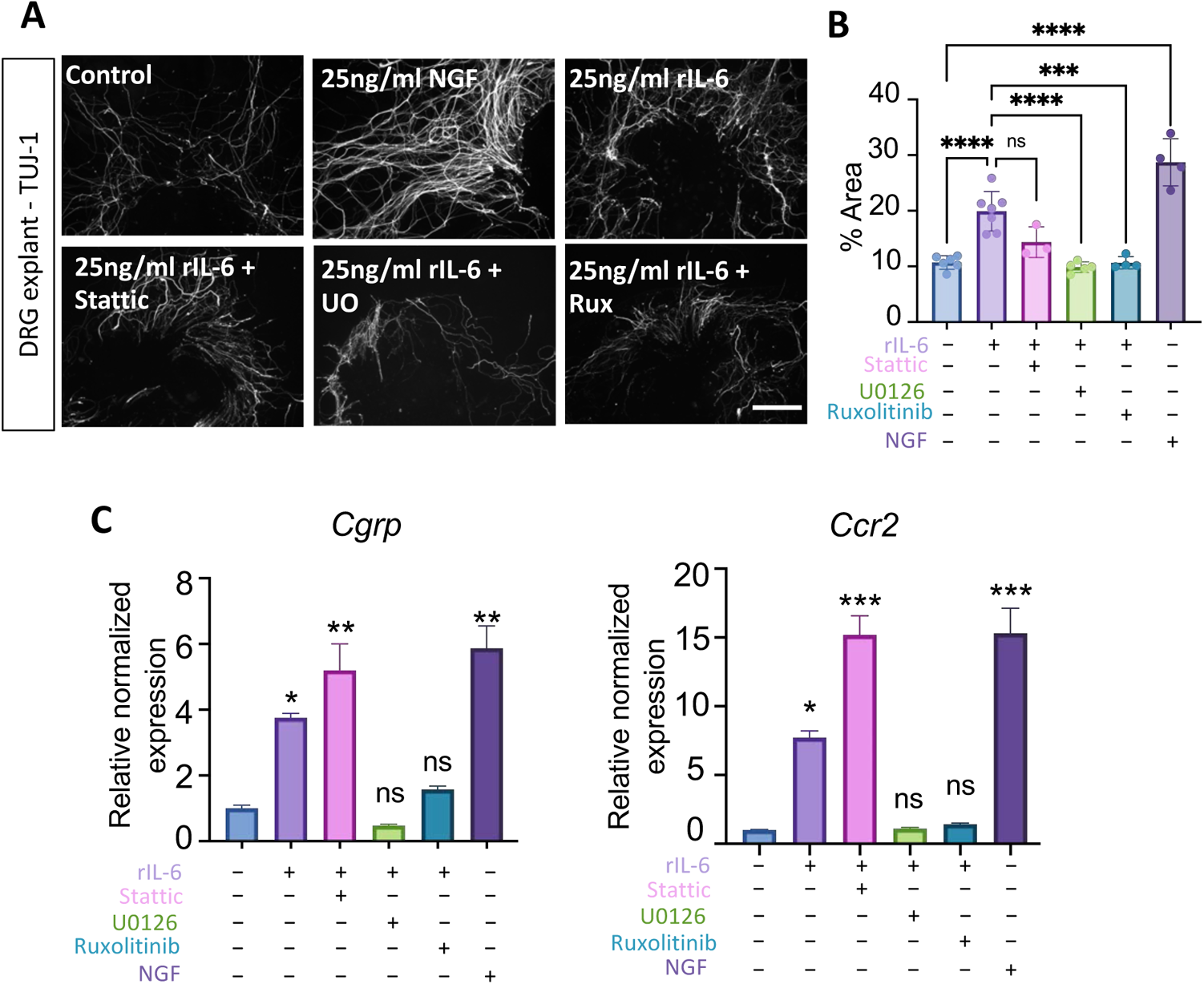
IL-6 induces pain-associated factors in DRG. (A) TUJ-1 staining of L3-L5 DRG cultured for 10 days with rIL6 and rNGF to demonstrate neurite outgrowth. N≥3 (B) Quantification of percentage of TUJ-1 staining area over the total examined area. Data represent mean ± SD. (**B**) qPCR of DRG neurons with DMSO or 50ng/ml IL-6 plus or minus inhibitors for 24 hours (10uM Ruxolitinib; 20uM Stattic; 10uM UO126). N≥ 3, p<0.05. Data represent mean ± SEM.* indicates statistical significance by one-way ANOVA between DMSO group and treatment group.

**Figure 7.**
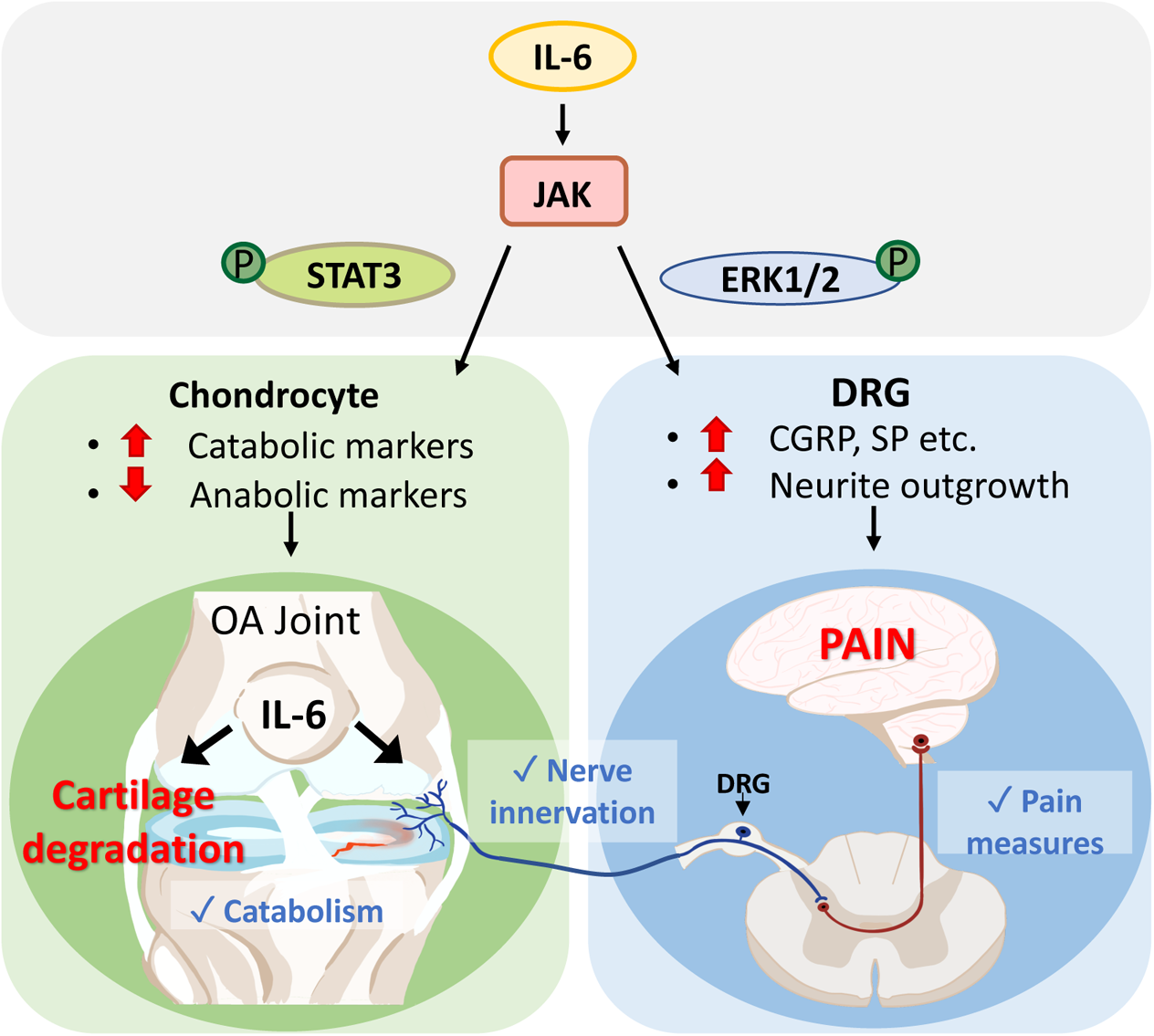
Model. A schematic model of this study

## Discussion

In this study, we identified a critical role for IL-6 signaling in mediating PTOA-associated cartilage degeneration and pain in males. Genetic removal of *Il6* reduced cartilage degradation via the attenuation of cartilage catabolism without affecting chondrocyte cell death. Additionally, deletion of *Il6* reduced PTOA-associated pain responses likely via the decrease in knee joint innervation and a reduction in pain mediators within knee-associated DRG neurons following joint injury. We further demonstrated the differential effects that downstream mediators of IL-6 signaling play in both chondrocytes and DRG neurons. JAK signaling was identified as a critical mediator of IL-6 induced cartilage catabolism and pain signaling in nociceptive neurons. We also demonstrated that STAT3 signaling is a potent regulator of cartilage catabolism; however, when inhibited demonstrated negative effects on cartilage anabolism and a further induction of pain-associated signaling in DRG neurons, suggesting that STAT3 inhibitors may not be an appropriate or effective DMOAD. ERK signaling was deemed to be essential for IL-6 induced neurite outgrowth and pain signaling in DRG neurons; however, inhibition of ERK1/2 only exhibited mild effects on cartilage catabolism. Collectively, this study provides novel insights into IL-6 and its downstream signaling effectors as potential therapeutic targets in the treatment of PTOA-associated cartilage degeneration and pain.

Of note, many of the conclusions from this study are based on the *in vivo* analysis of PTOA in male mice. Female mice are in general more refractory to PTOA-associated cartilage degradation than male mice^41^, although both exhibited similar pain responses following DMM injury as measured by the PAM device. Importantly, genetic removal of *Il6* in female mice did not alter the course of either PTOA-associated cartilage degeneration or pain over the 12-week assessments as observed in the *Il6* deficient male mice. These findings bring to the forefront two important points: 1) PTOA-associated chronic pain may be discordant with severe cartilage degeneration, since chronic pain is observed in both male and female mice at timepoints (early and late) in which visible cartilage effects of PTOA are extremely minor or absent, and 2) IL-6 signaling inhibition for the prevention of PTOA-associated cartilage degeneration and pain is only effective in males, which suggests that the underlying mechanisms regulating PTOA phenotypic progression may differ between males and females at the molecular level (at least in mice). Prior work in murine and cell models have demonstrated that estrogen is a negative regulator of *Il6*^44^, and that higher levels of estrogen receptor results in reduced *Il6* expression^45^. Since females exhibit higher levels of estrogen and estrogen receptor expression, females in general may have a more blunted effect to IL-6 signaling activation, such as the decreased severity of joint cartilage degradation following DMM injury observed in females as compared to male mice. Alternatively, females may have developed alternate pain signaling mechanisms to overcome the reduced effects of IL-6 signaling. Further investigations are still needed to identify the detailed gender differences in response to IL-6 signaling, IL-6 modulatory drugs, and their effects on PTOA-associated cartilage degeneration and pain.

IL-6 signaling modulators have been applied to the treatment of arthritis conditions in recent years. Tocilizumab, an antibody against the IL-6 receptor, is effective in treating rheumatoid arthritis (RA), and entered the market as a key disease-modifying anti-rheumatic drug (DMARD). As there are significant differences in the pathological mechanisms between RA and OA, the clinical results for their use in RA do not directly transfer to their use in OA conditions. A recent double-blinded, randomized controlled trial (NCT02477059) demonstrated that tocilizumab did not significantly reduce pain or improve function in patients with hand OA, and subsequently generated more adverse events such as infections and neutropenia within the treatment group^46^. This result is somewhat surprising in lieu of our findings, however hand OA is likely under the control of different signaling mechanisms as compared to PTOA of the knee. Further, signaling differences are known to exist between the membrane bound and soluble form of the IL-6 receptor, and therefore targeting generally the IL-6 receptor may have unintended consequences. Thus, a more specific downstream effector of the IL-6 receptor may be an even more appealing drug target providing better effectiveness with fewer side effects. Currently, small molecule JAK inhibitors are becoming the newest class of investigational drugs used to treat RA, and have proven efficacious in early pain relief and limiting the progression of structural damage in RA^47^. However, their use and effectiveness in OA or PTOA is unknown. Our study provides novel insights into the potential importance and application of JAK inhibition and their potential effects on cartilage degradation and pain in PTOA.

Our data demonstrate that STAT3 and ERK signaling regulate specific aspects of cartilage degradation and pain downstream of IL-6 signaling. Although studies have shown the STAT3 inhibitor, Stattic, to be effective in alleviating cartilage degradation in DMM-induced OA^32^, our data indicates that STAT3 inhibition may exacerbate IL-6 induced suppression of cartilage anabolic genes and further potentiate the expression of pain mediators. Therefore, drugs solely targeting STAT3 inhibition may not be the most all around effective therapeutic approach for treating PTOA-associated cartilage degeneration and pain. In opposition to STAT3 inhibition, ERK1/2 inhibition utilizing the drug, U0126, significantly reduced the expression of pain mediators induced by IL-6 signaling, however had limited efficacy in reducing IL-6 mediated cartilage catabolism. Therefore, the most effective approaches to treating both PTOA-associated cartilage catabolism and pain may lie in the dual inhibition of STAT3 and ERK signaling simultaneously, or alternatively the use of JAK inhibitors.

In conclusion, IL-6 signaling mediates both cartilage degradation and pain in the PTOA of males, but not females. Our study further defined the differential effects of downstream mediators of IL-6 signaling in both chondrocytes and DRG neurons, which has provided critical insights into not only the most relevant therapeutic targets of DMOADs, but also the population that would stand most to benefit from their use.

## Materials and Methods

### Mice

Interleukin-6 knockout mice (IL-6KO or *Il6^−/−^*) in the C57BL/6J background were used in this study and obtained from The Jackson Laboratory. C57BL/6J mice were used as wildtype (WT) controls and also obtained from The Jackson Laboratory. All surgeries were performed at 4 months of age. Mice were harvested at 4-weeks (5-month-old), 8-weeks (6-month-old) and 12-weeks (7-month-old) post injury. The numbers of mice used in each experiment are indicated in the Supplementary Table 1. All animal experiments were conducted in accordance with policies and regulations of the NIH and approved by the Duke University Institutional Animal Care and Use Committee (IACUC).

### Modified destabilized medial meniscus (DMM) OA model and surgical procedures

Destabilization of the medial meniscus was used as our model of post-traumatic OA (PTOA) in mice. We followed the well-established surgical method of destabilized medial meniscus with a slight modification.^29, 48, 49^ Briefly, the left hind-limbs were shaved and wiped with iodine and 70% ethanol to sanitize. After exposing thee left knee, the medial collateral ligament was transected. Next, the medial meniscus was destabilized from its anterior-medial tibial attachment using a 25G needle, and put back to its original location. The skin incision was sutured with 5-0 monofilament Nylon after the surgery. Sham surgeries were performed on separate animals with incisions made on the skin to expose the knee, however neither the ligament transection nor the meniscus detachment were performed.

### Histology

Knees were collected at 4-, 8- and 12-weeks post injury, and fixed with 4% PFA for two days at 4°C. Samples were decalcified with 14% EDTA for 10-14 days, and processed for paraffin embedding. The medial compartment of the knee joints were sectioned at 5-micron thickness.

Sections were collected at the point of meniscus separation, and stopped when the meniscus were no longer present in tissue sections, which resulted in about 60 sections per sample. We then performed immunohistochemistry (IHC) or immunofluorescence (IF), and evaluated cartilage degradation and other morphological changes by Safranin O staining with fast green counterstaining by following established protocols^50^. Specifically, sections were de-paraffinized, and unmasked for antigen with EDTA (pH =8) for phospho-STAT3 and IL-6 IHC, and citrate buffer for phospho-ERK1/2 IHC at 95°C. MMP-13 IF staining were performed with paraffin slides as described previously^51^. Cell apoptosis was examined utilizing a TUNEL (Terminal deoxynucleotidyl transferase dUTP nick end labeling) kit (Roche). In brief, after de-paraffinization, sections were subjected to antigen retrieval with proteinase K (10ug/ml) for 20 mins at room temperature. After washes with PBS, sections were incubated in a humidified chamber with the TUNEL reaction mix for 60 minutes at 37°C, followed by coverslipping and imaging.

In addition to knee joint tissues, DRG at L3-L5 were harvested at the same time, and RNA, protein, neuronal cultures, and frozen sections were prepared sequentially from these DRG harvests. DRG were snap frozen in liquid nitrogen and stored at −80 degrees for RNA and protein extraction. For frozen sections, DRGs were harvested after transcardial perfusion of mice and processed through a gradient of sucrose from 15% to 30% for 2 days before embedding in O.C.T. DRG explant cultures were adapted from established protocols^52^. For TUJ-1 staining on DRG explants, DRGs were fix with 3.7% formaldehyde for 30 minutes, and then incubated with TUJ-1 antibody as a 1:2000 dilution in PBS buffer with 0.3% Triton-100 and 10% normal goat serum overnight at 4°C. After incubation with appropriate secondary antibody diluted in PBS, sections were coverslipped with DAPI and prepared for imaging.

### Osteoarthritis Research Society International (OARSI) osteoarthritis cartilage histopathology assessment (OARSI Score)

OARSI scoring is a standard histological scoring system for murine OA^53, 54^. In our experiments, five sections at different depths, approximately 50um apart, were selected for each animal. Based on Safranin-O stained sections, cartilage changes were scored blindly from 0 to 6 by at least three individuals. Every data point in the OARSI score panels represent one animal, which is an average score from three graders on five individual sections per animal. A score of 0-2, represents only minor changes observed within the articular cartilage, while scores of 3-6 indicate increasing areas of cartilage erosion across the articular surface.

### MicroCT analysis

MicroCT (uCT) analysis were performed on knees from WT control and *Il6^−/−^* mice prior to decalcification using a VivaCT 80 scanner with 55-kVp source (Scanco USA) as previously described^55^. Quantification of microCT data were calculated for the subchondral region, which we defined as 0.1mm region above proximal growth plate of the tibia extending to the articular cartilage. Bone volume/total volume (BV/TV) at the region of interest is used as primary outcome parameter.

### Behavioral testing

For pain measurements, behavioral tests were conducted before harvesting tissue at 1-, 2-, 4-, 6-, and 8-weeks post-injury (Sham/DMM) using the Pressure Application Measurement (PAM) device to assess mechanical hyperalgesia^56–59^. Animals were adapted to the test environment with stable room temperature and humidity for two days before the baseline testing. In PAM tests, a gradually increasing force was applied across the knee joint of the mouse which is lightly but securely held, until the animal provides an indication of pain or discomfort. The peak force applied within the 5 second maximum test duration is recorded by the device connecting to force sensor worn on the operator’s finger and is displayed in grams as the paw withdrawal threshold.

### Cell and Ex vivo cartilage and DRG explant cultures

ATDC5 cells (RIKEN BioResource Center) were maintained and cultured as previously described^31^. In short, ATDC5 cells were maintained in complete media DMEM/F-12 (1:1) medium with addition of 10% FBS and 1% Penicillin/Streptomycin. For experimental purposes, ATDC5 cells were differentiated in complete media supplemented with 0.1% insulin, transferrin, and sodium selenite (ITS) for 7 to 14-days. Before treatment with rIL-6 protein or inhibitors, differentiated ATDC5 chondrogenic cells were washed with PBS once, and the indicated treatments were added to the media. The JAK inhibitor, Ruxolitinib (Selleckchem), was used at a final concentration of 0.5uM; the STAT3 inhibitor, Stattic (Sigma), was used at a final concentration of 20uM; and the ERK1/2 inhibitor, U0126 (Sigma), was used at a final concentration of 10uM.

For cartilage explant culture, femoral cartilages were harvested from three-week-old mice and cultured in same media as ATDC5 cell (DMEM/F-12 medium + 10% FBS + 1% Penicillin/Streptomycin + ITS) with 500ng/ml recombinant IL-6 (rIL-6) protein for 7 to 10-days. DRG explant cultures are adapted from the protocol described previously^52^. Briefly, following DRG harvest from 1 month old WT mice, the DRGs were placed in a 35mm Petri dish containing 3 mL of ice-cold serum free Neurobasal-A media with 1% Penicillin/Streptomycin. Each individual DRG was cleaned and trimmed of excess fibers and connective tissues under a surgical microscope and was placed in a new petri dish containing ice-cold media. All steps after this were performed using aseptic technique in a Class II biosafety cabinet. DRG were washed with sterile HBSS solution twice and plated in PDL pre-coated plates. Neurobasal medium supplemented with 2% v/v B-27 Supplement were gently added to cover and maintain the entire explant at 37°C and 5% CO_2_ in a cell culture incubator.

### Gene and protein analysis

ATDC5 cells and DRGs were collected in RLT buffer for RNA extraction and RIPA lysis buffer (radioimmunoprecipitation assay (RIPA) buffer plus protease inhibitor and phosphatase inhibitors) for protein extraction. A motorized tissue grinder was used to homogenize DRG in order to extract RNA or protein. RNA was extracted with the RNeasy Mini Kit (Qiagen) according to the manufacturer’s instructions, and RNA concentrations were measured using a NanoDrop 1000 spectrophotometer. After RNA extraction, cDNA were synthesized and real-time qPCR analysis were performed for the chondrogenic related genes (*Acan, Col2a1, MMP-13, Adamts5*), several pain mediators (*Cgrp, and Ccr2*) and a housekeeping gene, *Gapdh* (see Supplementary Table 2 for sequences). For Western analysis, 30ug of protein were loaded to each well, and underwent electrophoresis through 4-20% gradient Tris-Glycine gels. Proteins were transferred to polyvinylidene difluoride (PVDF) membranes with a Trans-Blot Turbo Transfer System (Bio-Rad). IL-6 downstream signaling activation was monitored using primary antibodies against STAT3, pSTAT3 (Tyr705), ERK1/2, pERK1/2 (Thr202/Tyr204), p38 MAPK, and pp38 MAPK (Thr180/Tyr182), which were incubated overnight in 1:1000 dilution in 5% nonfat milk dissolved in 1x TBST. After blotting with appropriate secondary antibodies, gels were developed using the Pierce ECL Western Blotting Substrate or SuperSignal West Femto Maximum Sensitivity Substrate depending on the intensity and were imaged using a Chemidoc system (Bio-Rad).

### Statistical analysis

Statistical significance and p-values were determined using one-way ANOVA for OARSI scores, quantification of MMP-13 and TUNEL IF stainings, and BV/TV values from our uCT analyses. Two-way ANOVA with Bonferroni’s post-hoc test were used for behavioral analysis, and unpaired two-tailed t-tests were utilized for qPCR results (See Supplementary Table 1).

## List of Supplementary Materials

Supplementary Fig. 1. DMM injury induces progressive PTOA phenotypes

Supplementary Fig. 2. No differences between *WT* and *Il6−/−* mice within knee joints and DRG

Supplementary Fig. 3. Articular chondrocyte apoptosis is not reduced via the loss of IL-6 during PTOA development.

Supplementary Fig.4. Cartilage and pain phenotypes in female *WT* and *Il6−/−* mice with PTOA

Supplementary Fig. 5. Culture models and validation of pathway inhibitors

Supplementary Fig. 6. Neurite outgrowth in isolated DRG neurons

Supplementary Table 1. Sample sizes and statistics

Supplementary Table 2. qPCR primer sequences

Supplementary Table 3. Mice and reagents

## Acknowledgements

We would like to thank Dr. Yun Gu, Dr. Deepika Sharma, and the Duke Light Microscopy Core Facility for technical and equipment support.

## Author Contributions

Y.L: Conceptualization, Methodology, Investigation, Formal Analysis, Writing – Original Draft, Review, and Editing, Visualization. X.L: Investigation, Formal Analysis, Writing – Review and Editing. Y.R: Investigation, Formal Analysis, Writing – Review and Editing. J.T.L: Investigation, Writing – Review and Editing. A.J.M: Methodology, Writing – Review and Editing, A.P.L: Investigation, Writing – Review and Editing. R-R.J: Supervision, Resources, Writing – Review and Editing. M.J.H: Conceptualization, Methodology, Writing – Review, and Editing, Visualization, Supervision, Project Administration, and Funding Acquisition.

## Data and Material Availability

All data associated with this study are available in the main text or the supplementary materials.

## Supplementary Figures and Legends

**Supplementary Figure 1.**
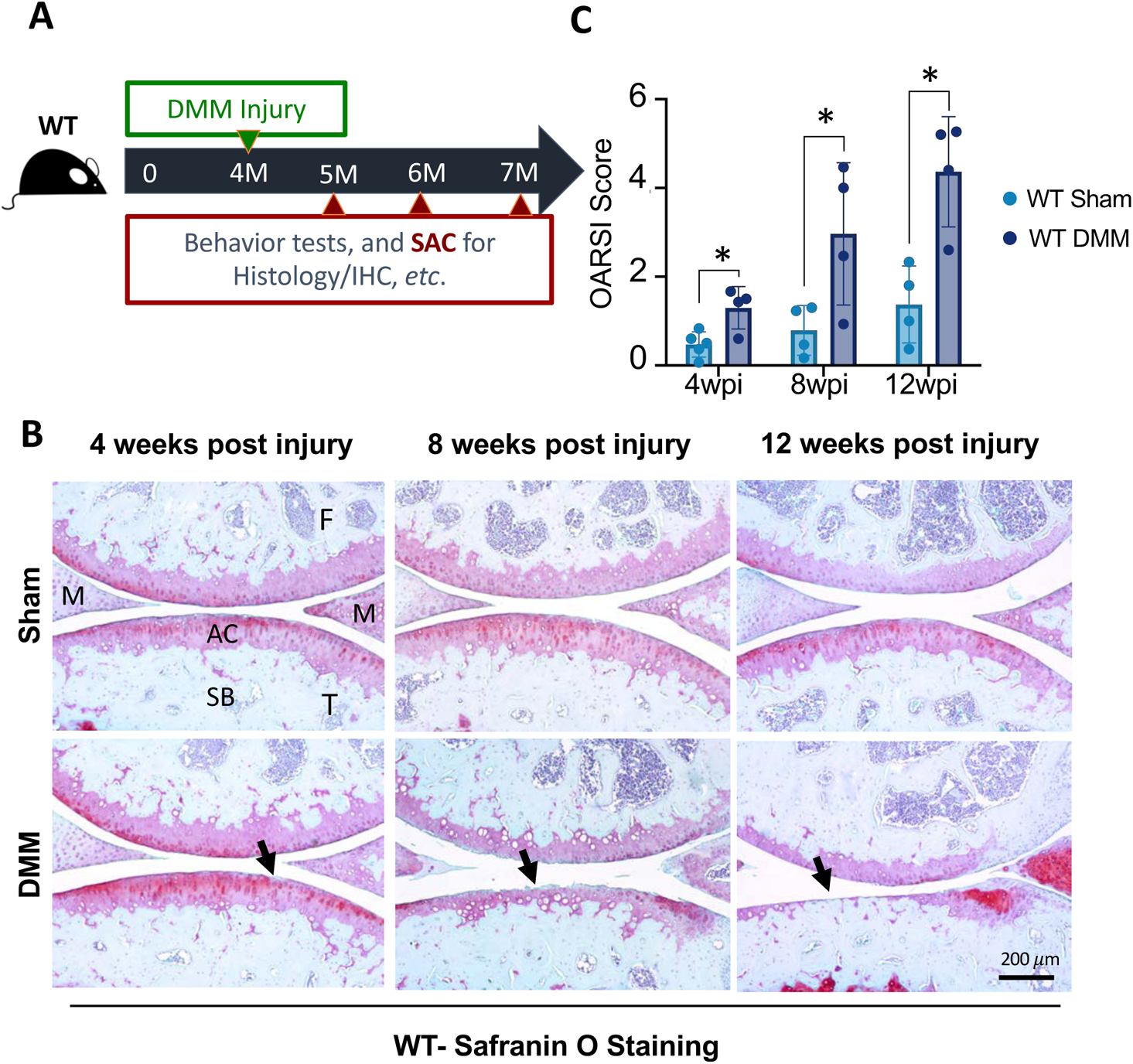
DMM injury induces progressive PTOA phenotypes. (A) A model showing the schematic of DMM/sham surgery in WT mice (B) Safranin-O staining of WT knee joint sections at 4-, 8- and 12- weeks post injury in male mice. (C) OARSI scores of knee sections from male mice at 4- (N≥4), 8- (N=4) and 12- (N=4) weeks post injury. p<0.05; One-way ANOVA. Bar = 200 μm. Data represent mean ± SD.

**Supplementary Figure 2.**
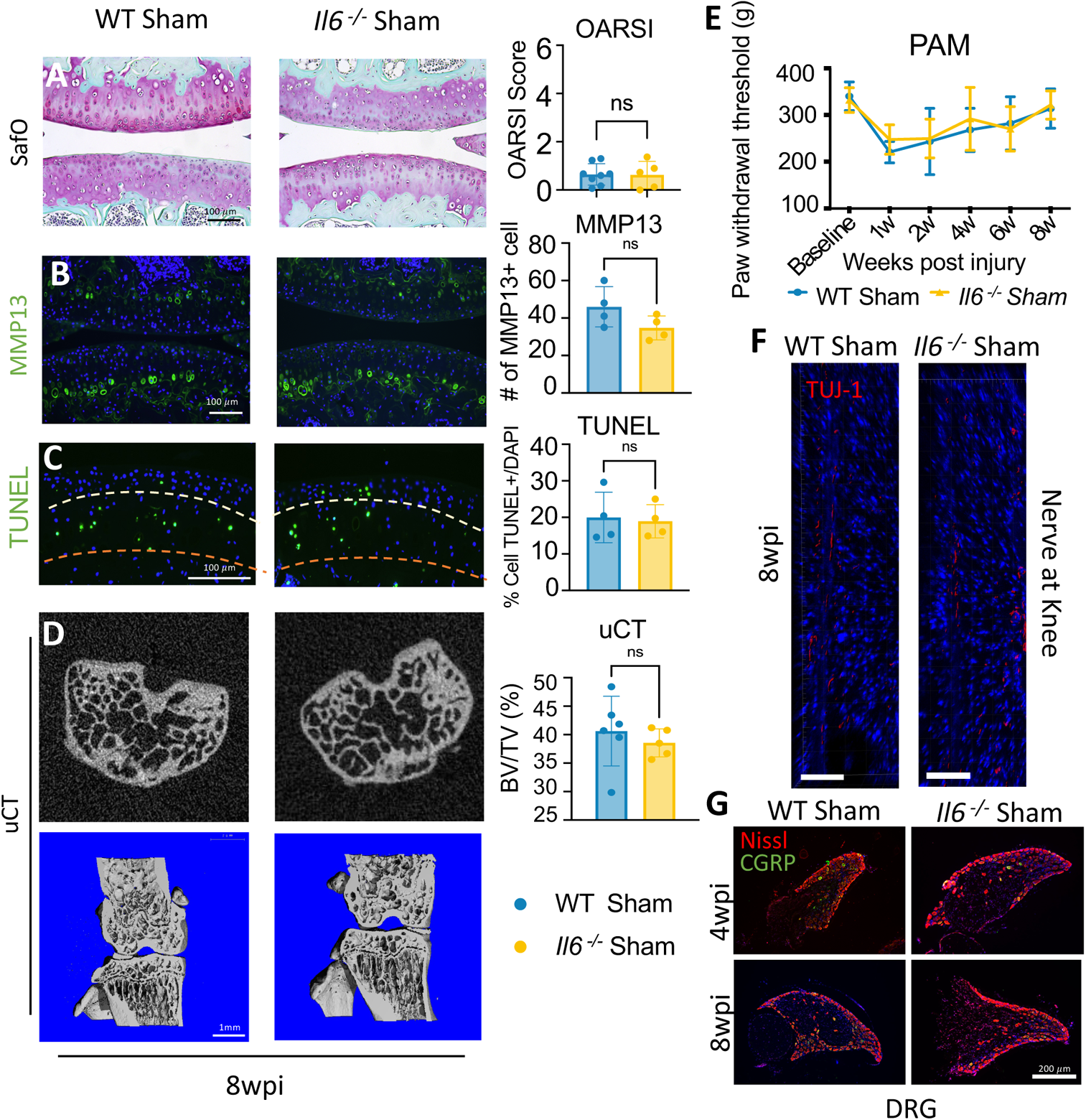
No differences between *WT* and *Il6−/−* mice within knee joints and DRG. (A) Safranin-O staining of WT and *Il6−/−* knee joint sections at 8-weeks post sham injury in male mice (N≥ 5); Bar = 100 μm. On the right, OARSI score quantified the cartilage phenotype at 8-weeks post sham injury (B) IF staining of MMP-13 (green) of knee sections from WT or *Il6−/−* mice at 8-weeks post sham surgery (N≥4); Bar = 100 μm. Quantification of the number of cell with MMP-13 staining on the right. (C) TUNEL staining in WT and *Il6−/−* knee joint sections at 8-weeks post sham injury in male mice. (N≥ 4); Bar = 100 μm. Quantification of TUNEL staining on the right. (D) MicroCT analysis shows cross and sagittal images of the knee of WT or *Il6−/−* mice 8- weeks post sham injury. (N≥5); Bar = 1 mm. Quantification of subchondral bone with 0.1mm thickness of the region of interest is evaluated on the right side. (E) PAM measurements of knee pain in WT or *Il6−/−* male mice with sham injury (N≥ 8). (F) Characterization of TUJ-1+ innervation at the knee joint of WT or *Il6−/−* male mice 8-weeks post sham injury (N≥ 3) (G) IF of CGRP+ neuron in L3-5 DRG from WT or *Il6−/−* mice at 4-and 8- weeks post sham injury. CGRP in green, Nissl in red (N≥ 4); Bar = 200 μm. All statistics were unpaired two-tailed t-test; p<0.05. Data represent mean ± SD.

**Supplementary Figure 3.**
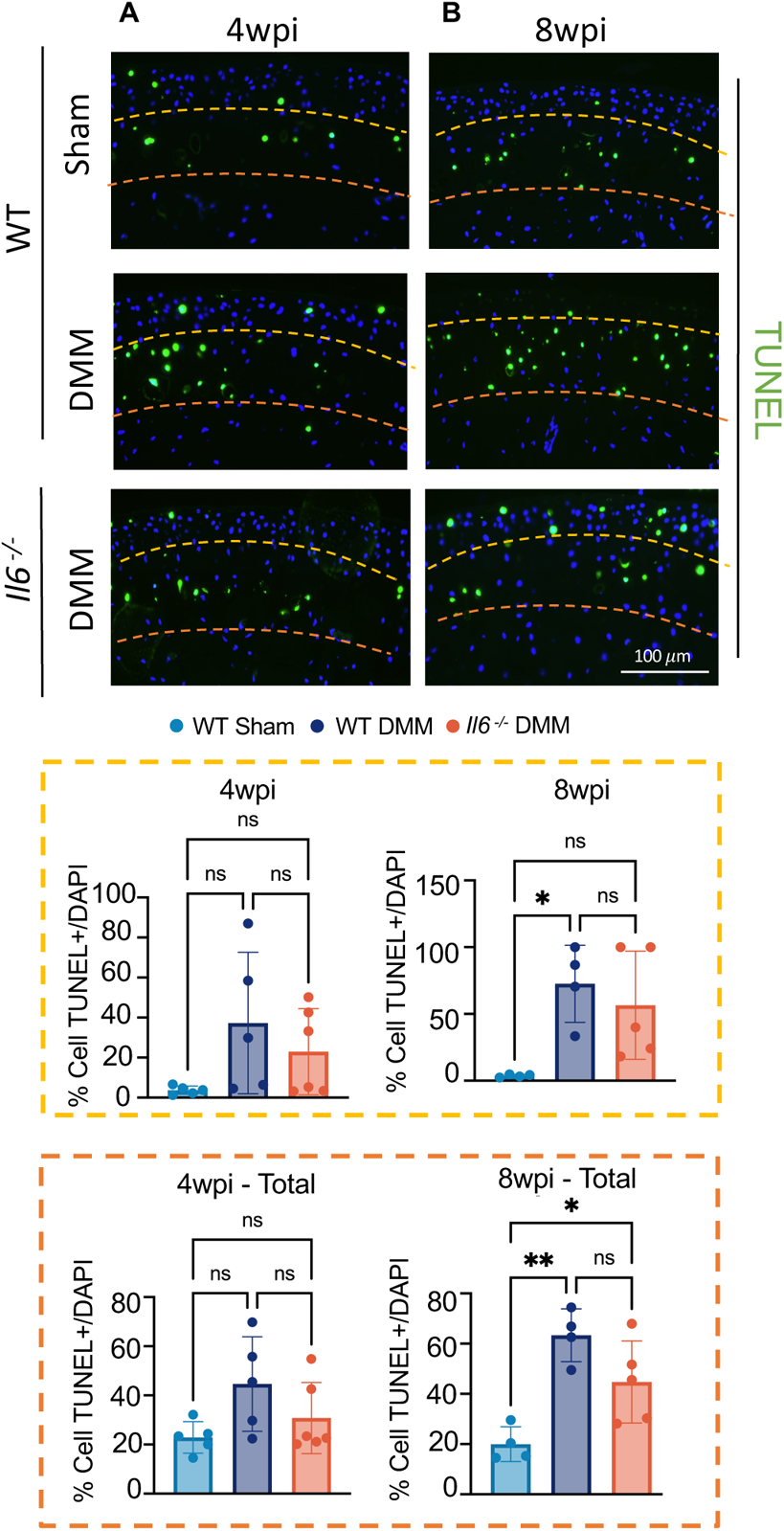
Articular chondrocyte apoptosis is not reduced via the loss of IL- 6 during PTOA development. TUNEL staining (green) of knee sections from WT or *Il6−/−* mice at 4- (A), and 8- (B) weeks post sham or DMM surgery. Percentage of TUNEL+ cells to total cells above tidemark (marked in yellow line); Percentage of TUNEL+ cells to total cells above calcified region (marked in orange line) N≥4); Bar = 100 μm. p<0.05; One-way ANOVA. Data represent mean ± SD.

**Supplementary Figure 4.**
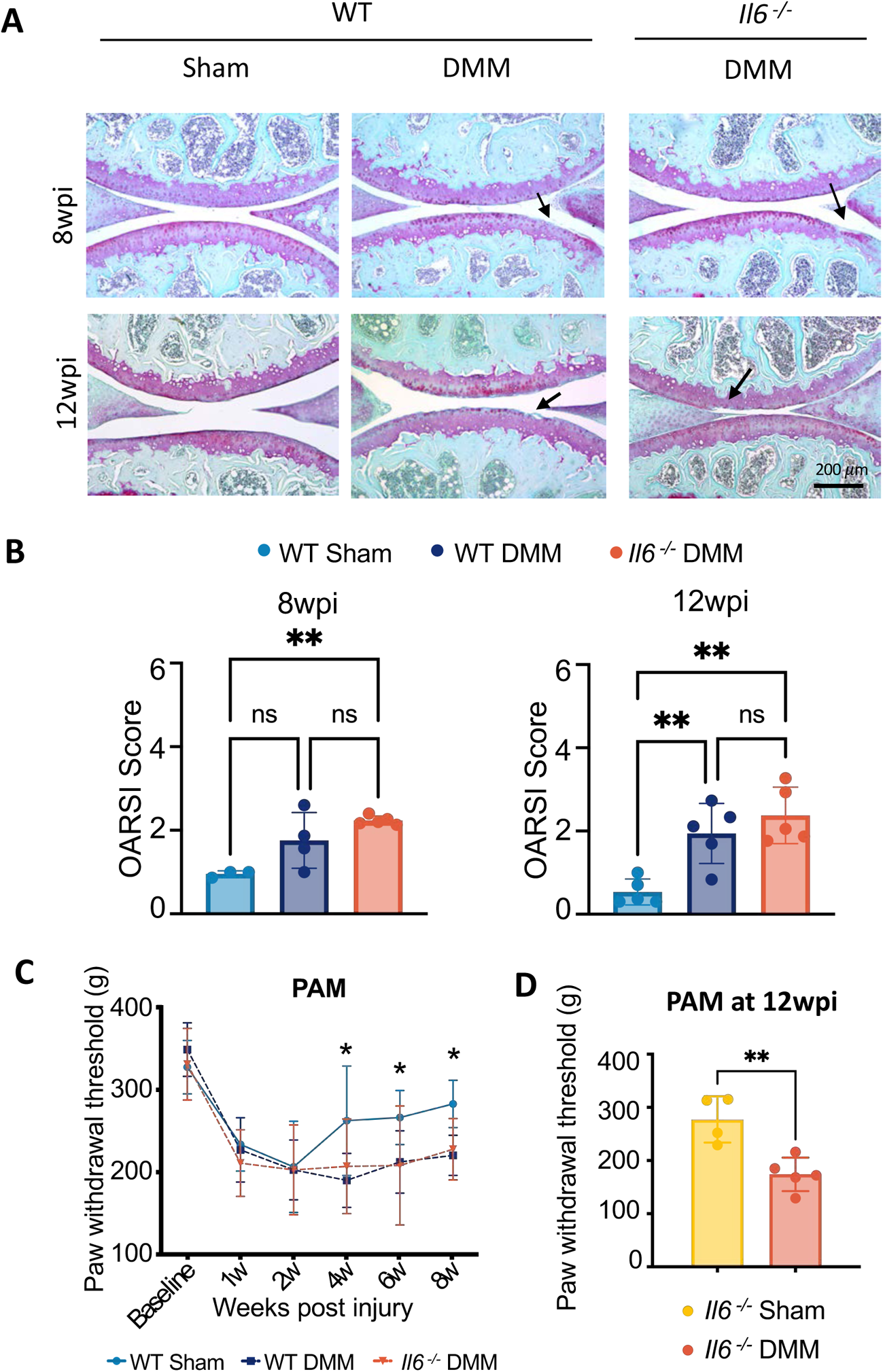
Cartilage and pain phenotypes in female *WT* and *Il6−/−* mice with PTOA. (A) Safranin-O staining of WT and *Il6−/−* knee joint sections at 8- and 12-weeks after injury in female mice. N≥3; Bar = 200 μm. (B) OARSI scores at 8- (N≥ 3), and 12- (N≥ 5) week post injury. p<0.05; One-way ANOVA. Data represent mean ± SD. (C) PAM measures of knee pain in female mice from 1- to 8-weeks post injury. N≥ 8 p<0.05; Two-Way ANOVA with Bonferroni’s post-hoc test. (D) PAM assessment of *Il6−/−* knee pain at 12 weeks post injury. N≥ 4 p<0.05; One-way ANOVA. Data represent mean ± SD.

**Supplementary Figure 5.**
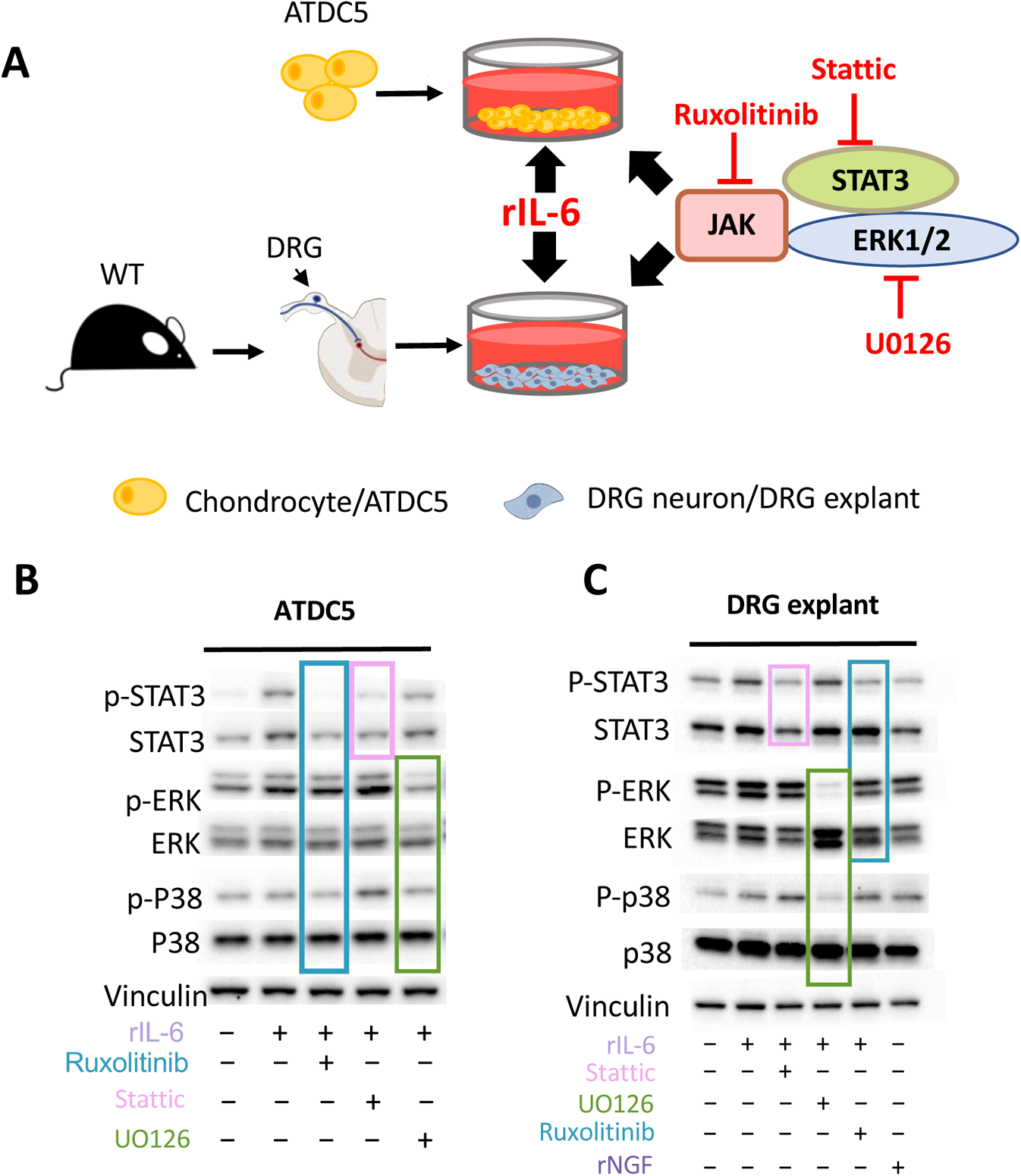
Culture models and validation of pathway inhibitors. (A) An illustration of cell culture systems utilized for Figure 5 and 6 (B) Western blot on protein from ATDC5 cultures with treatment of rIL-6 and specified inhibitors (0.5uM Ruxolitinib; 20uM Stattic; 10uM U0126). 30 minute treatments. N≥ 3 (C) Western blot shows the efficiency of the inhibitors in DRG explant culture (10uM Ruxolitinib; 20uM Stattic; 10uM U0126). N≥ 3.

**Supplementary Figure 6.**
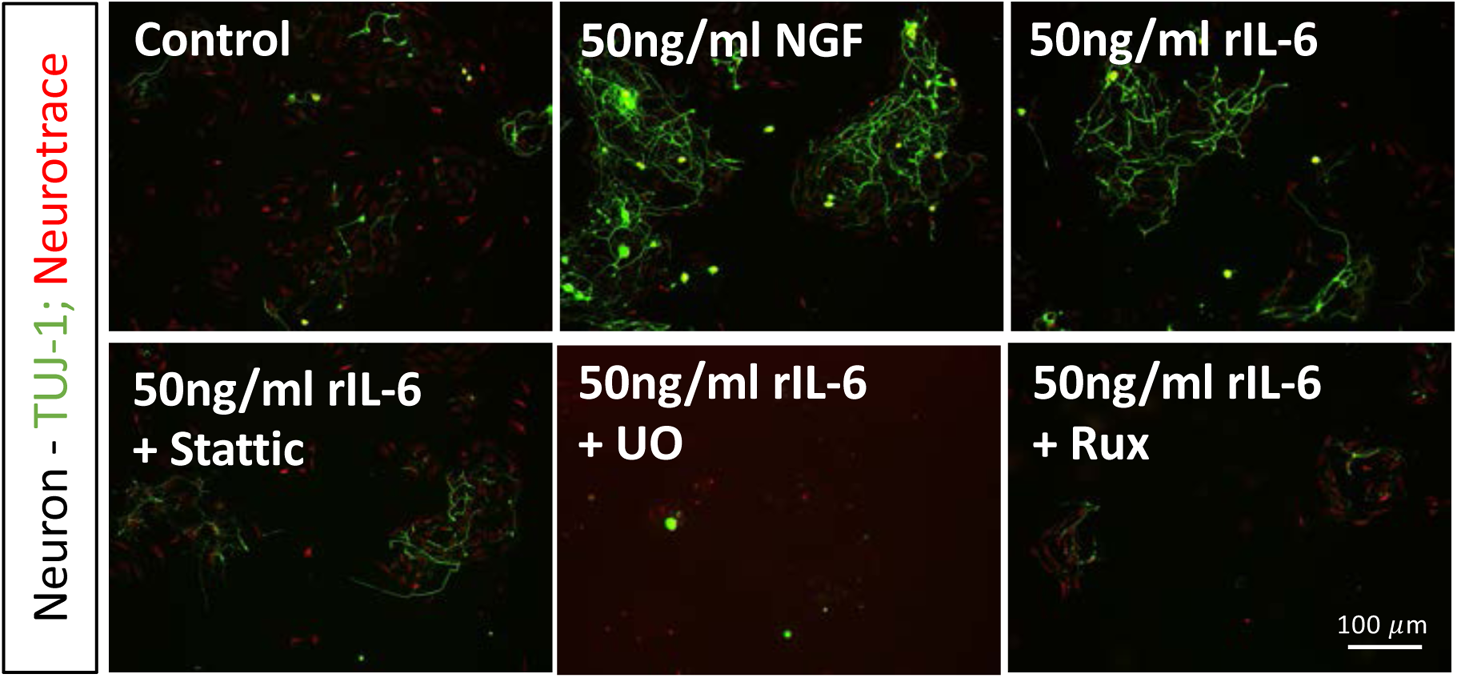
Neurite outgrowth in isolated DRG neurons. TUJ-1 (Green) staining of isolated primary neurons from L3-L5 DRG cultured for 10 days with DMSO or 50ng/ml rIL-6 plus or minus inhibitors for 24 hours (10uM Ruxolitinib; 20uM Stattic; 10uM U0126). 50ng/ml recombinant NGF were used as a positive control. N≥3; Bar = 100 μm.

